# Mean-field theory accurately captures the variation of copy number distributions across the mRNA’s life cycle

**DOI:** 10.1101/2021.08.24.457469

**Authors:** Juraj Szavits-Nossan, Ramon Grima

## Abstract

We consider a stochastic model where a gene switches between two states, an mRNA transcript is released in the active state and subsequently it undergoes an arbitrary number of sequential unimolecular steps before being degraded. The reactions effectively describe various stages of the mRNA life cycle such as initiation, elongation, termination, splicing, export and degradation. We construct a novel mean-field approach that leads to closed-form steady-state distributions for the number of transcript molecules at each stage of the mRNA life cycle. By comparison with stochastic simulations, we show that the approximation is highly accurate over all of parameter space, independent of the type of expression (constitutive or bursty) and of the shape of the distribution (unimodal, bimodal and nearly bimodal). The theory predicts that in a population of identical cells, any bimodality is gradually washed away as the mRNA progresses through its life cycle.

## I. INTRODUCTION

In recent years, the study of stochastic models of gene expression has received wide attention [1]. In the absence of transcriptional feedback, these models are often composed of first-order reactions, in which case exact expressions can be derived from the master equation for all the moments of the distribution of mRNA numbers. This is because for linear propensities, the moment equations are closed and can be solved straightforwardly [2].

However the exact closed-form solution of the master equation for the probability distribution of mRNA numbers is a much harder problem. Popular methods such as the Poisson representation of Gardiner [3] and those devised more recently by Jahnke and Huisanga [4] are not applicable to models of gene expression composed of only first-order reactions because of the presence of a reaction of the type *G* → *G* + *M* which models transcription of mRNA (*M*) from the active state of a promoter (*G*). Hence the general solution of these models is still an open research question. However progress has been achieved on the solution of specific models.

A two-state model (commonly known as the telegraph model) whereby a gene switches between two states (an active and an inactive state) and mRNA is transcribed in the active state, has been solved exactly in steady-state [5] and in time [6] for the marginal distribution of mRNA numbers. Various other extensions of this model to include more biological realism have also been solved exactly or approximately for the marginal distribution of mRNA numbers. These include models that account for more than two gene states [7–9], for polymerase dynamics [10], for leaky expression from the inactive state [11], cell-to-cell variability (static and dynamic) [11, 12], replication and binomial partitioning due to cell division [13], cell cycle duration variability [14] and modulation under environmental changes [15].

We here consider a different type of extension of the telegraph model that has recently received attention in three different biological contexts: (i) RNA polymerase (RNAP) movement along a gene during transcription [16]; (ii) multi-step splicing [17]; (iii) nuclear retention of mRNA [18]. In this stochastic model, a gene switches between two states (an active state *G* and an inactive one *G*^∗^), produces a transcript in the active state and subsequently it undergoes an arbitrary number *L* of sequential unimolecular steps, leading to *L* forms of the transcript (denoted by *P*_*i*_, where *i* ∈ [1, *L*]). Each transcript either is removed after the *L* steps or else can also be degraded prematurely at an earlier step. A reaction scheme for this model is as follows:

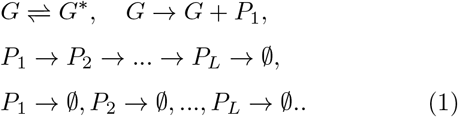

The *L* downstream processing steps can be interpreted in various ways. At the coarsest scale, the model with *L* = 2 steps can capture nuclear mRNA and cytoplasmic mRNA dynamics. At a less coarser scale, the model can capture the dynamics of the whole mRNA life cycle from birth (initiation) to death (in the cytoplasm). For example one can associate steps *i* = 1, …, *L*_1_ − 1 with elongation of the nascent transcript, *i* = *L*_1_ with termination, *i* = *L*_1_ + 1, …, *L*_2_ − 1 with splicing, *i* = *L*_2_ with export from the nucleus to the cytoplasm and *i* = *L*_2_ + 1, …, *L* with several reaction steps leading to mRNA degradation. For the latter process, for example, the sequence of steps can model the fact that the polyadenylate tails of eukaryotic transcripts are sequentially chewed up before the protein-coding part of the message is degraded [19]. At a fine scale, the model could capture the dynamics of nascent mRNA only, where the *L* steps represent the unidirectional movement of RNAP (with a nascent mRNA tail attached to it) along the gene. Hence the reaction scheme (1) has myriad molecular interpretations according to the desired application.

An exact, steady-state and closed-form solution for the marginal distribution of the number of molecules of *P*_*i*_ is unknown. In this paper, we report an approximate solution to this open problem. Because the method we devise is non-perturbative and does not make assumptions on the strength of correlations between the forms of the product, we find by comparison with stochastic simulations that the approximation is highly accurately across all of parameter space.

The paper is divided as follows. In Section II, we describe the model in detail, formulate its master equation and discuss some known exactly solvable cases. In Section III, we derive exact expressions for the moments of *P*_*i*_ using a lattice path method. In Section IV, we use the first and second moments and mean-field approximation to construct marginal distributions for *P*_*i*_. We show that while the use of standard mean-field approximation is valid in only certain regions of parameter space, use of a novel type of mean-field approximation leads to a uniformly accurate approximation over all parameter space. In particular, we show that the latter is practically indistinguishable from distributions calculated using the stochastic simulation algorithm. We conclude in Section V by a discussion of our method versus those already in the literature.

## II. STOCHASTIC MODEL OF TRANSCRIPTION KINETICS

In this Section we define the model and introduce the master equation that describes its stochastic evolution in time. While as mentioned in the Introduction, the model has many interpretations, in what follows we will describe it in terms of RNAP dynamics during transcription.

### A. Definition of the model

Transcription is modelled as a series of steps involving promoter activation, initiation at the transcription start site, RNAP movement along the gene and termination at the stop site that frees the newly synthesized mRNA.

The promoter is modelled having two states, activated (on state *G*) and deactivated (off state *G*^∗^). Gene body— the part of the gene between the start and stop sites— is divided into *L* segments. The state of the system is described by the state of the promoter (on or off), and the number of RNA polymerases in each of the *L* segments, *n*_*i*_ for *i* = 1, …, *L*.

The model is summarised in Fig. 1 – note that this is a cartoon version of the reaction scheme (1) which is specific to the context of transcription. In particular the RNAP on gene segment *i* is what was labelled as *P*_*i*_ in scheme (1). Note that the number of RNAPs on a gene is equal to the number of nascent mRNA; this is because each RNAP has attached to it a nascent mRNA tail whose length increases as elongation proceeds. The rates of gene activation and deactivation are *s*_*u*_ and *s*_*b*_, respectively. Once the promoter is active (in the on state), transcription initiation occurs at rate *r*, whereby a new RNAP enters the first segment. We do not consider leaky transcription, i.e. transcription initiation is not allowed when the promoter is in the off state. After initiation, the RNA polymerese moves along the gene body from segment to segment with, in general, segment-dependent elongation rate *k*_el,*i*_. During elongation, the RNAP can prematurely detach from the gene at rate *d*. Transcription termination occurs at the last segment with rate *k*_el,*L*_, whereby the RNAP is removed from the gene and the newly synthesized mRNA is released.

**FIG. 1.**
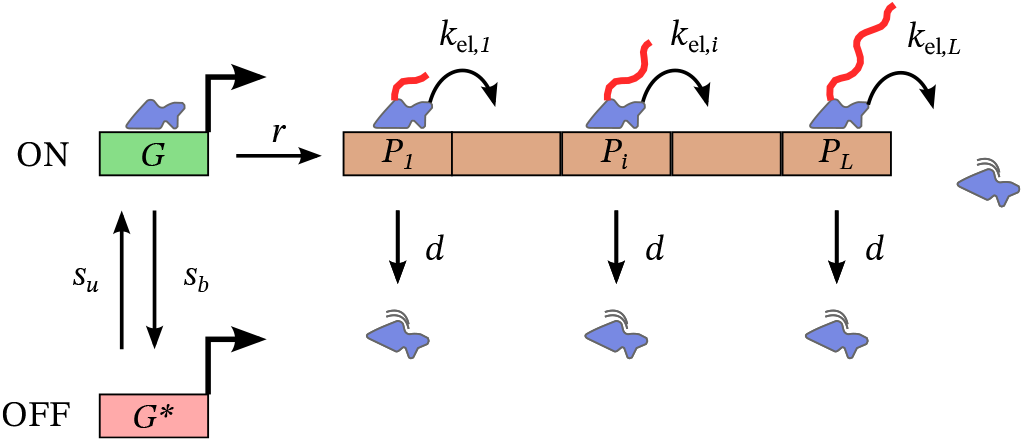
Schematic of the stochastic transcription model. The gene body is divided into *L* segments. The promoter can be in two states: active (on) and inactive (off). Promoter on and off switching rates are *s*_*u*_ and *s*_*b*_, respectively. Transcription initiation occurs when the promoter is in the on state at rate *r* by which a new RNAP is added to the first segment. No transcription initiation is allowed when the promoter is in the off state. Following initiation, RNAPs move along the gene with segment-dependent elongation rates *k*_el,*i*_. They can detach from DNA at any segment with rate *d* (premature termination). Once they reach the stop site they terminate transcription at rate *k*_el,*L*_. The model ignores excluded volume interactions between neighbouring RNAPs, i.e. the number of RNAPs in each segment is unbounded.

We note that the model does not explicitly take into account excluded volume interactions between RNAPs. RNAP has a footprint of *ℓ*_*RNAP*_ = 35 base pairs (bp) [20]. The size of each segment is *ℓ*_segment_ = *L*_gene_*/L*, where *L*_gene_ is the number of base pairs in the gene body, typically measured in thousands of base pairs. Thus the maximum number of RNAPs in each segment is *c* = *ℓ*_segment_*/ℓ*_RNAP_, whereas in our model the number of RNAPs in each segment is unbounded.

As we show later, this simplification allows us to obtain analytical expressions for the distribution of RNAPs along the gene. In order to compensate for this simplification, one can choose the size of the segment *ℓ*_segment_ such that the average number of RNAPs in any segment is less than the maximum capacity *c*

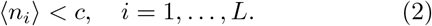

Alternatively, one can choose *ℓ*_segment_ such that the probability of observing more than *c* RNAPs in each segment is small,

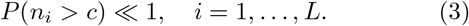

We provide analytical expression for both ⟨*n*_*i*_⟩ and *P*(*n*_*i*_) so that any of these two conditions can be easily checked if the model parameters are known. Transcription is typically rate-limited by initiation, which means there are only few active RNAPs on the gene at a given time, and hence conditions (2) and (3) are automatically satisfied.

### B. Master equations for the RNAP distribution

The central information that we are interested in is the probability *P* (*n*_1_, …, *n*_*L*_; *t*) to find *n*_1_ RNAPs in segment 1, *n*_2_ RNAPs in segment 2, and so on, at a given time *t*. This probability is in fact a sum of two probabilities *P*_on_(*n*_1_, …, *n*_*L*_; *t*) and *P*_off_ (*n*_1_, …, *n*_*L*_; *t*) which are conditioned on the state of the promoter. The probabilities *P*_on_ and *P*_off_ satisfy the following master equations,

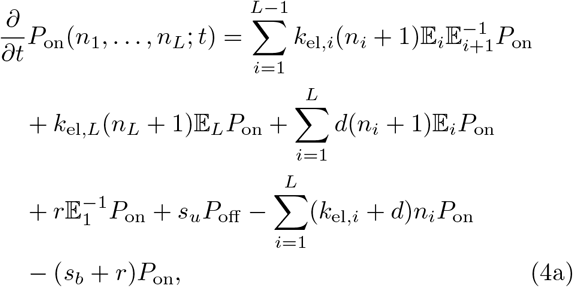

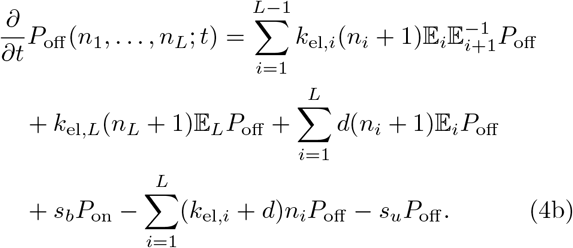

Here we have shortened the notation by introducing step operators 𝔼_*i*_ and 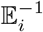 [21] that increase and decrease the number of particles *n*_*i*_ in segment *i*, respectively,

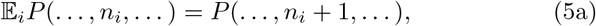

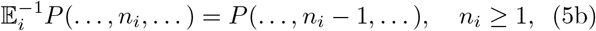

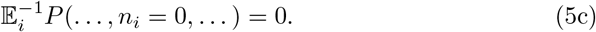

From now on, we are interested only in the steady state, so that *∂P*_on_*/∂t* = 0 and *∂P*_off_ */∂t* = 0 and so we drop the time dependence from *P*_on_ and *P*_off_.

Rather than working with the master equation directly, we introduce the generating functions *G*_on_, *G*_off_ for each of the promoter states, and also *G* = *G*_on_ + *G*_off_, whereby

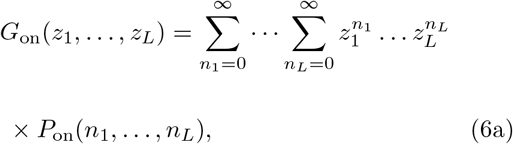

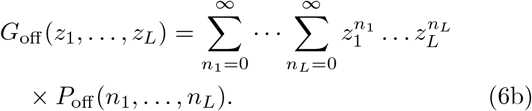

From the master equation we get the following steady-state partial differential equations for *G*_on_ and *G*_off_,

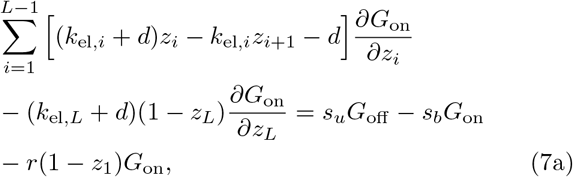

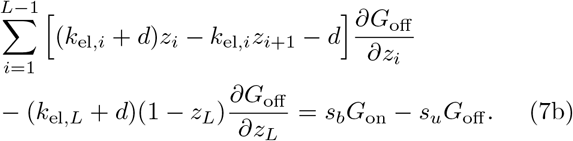

To the best of our knowledge, this system of equations has not been solved in closed-form before, except in three special cases: telegraph model (*L* = 1), constitutive gene expression (*s*_*b*_ = 0) and deterministic elongation with mature RNA production. For completeness we summarise these three cases below.

### C. Exactly solvable cases

#### 1. The telegraph model

The case of *L* = 1 is known as the telegraph model [5], whose solution for the generating function reads

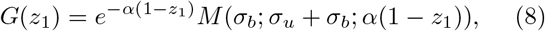

where *k*_el,1_ = *k*_el_ and

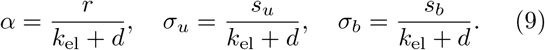

In the expression for *G*(*z*_1_), *M* is Kummer’s (confluent hypergeometric) function [22]. The probability distribution of *n*_1_ is given by

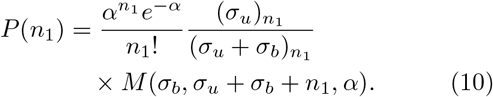

We note this expression also holds for the marginal distribution *P*(*n*_1_) of the model with *L* ≥ 1 – this is because the fluctuations in the RNAP numbers on segment 1 are unaffected by the fluctuations on the other segments due to the irreversible nature of the reactions between segments.

#### 2. Constitutive gene expression

A constitutive promoter is always in the on state, which means that *s*_*b*_ = 0. In that case there is only one equation to solve, that for *G*_on_ ≡ *G*, since *G*_off_ = 0. We can easily check that the solution is given by

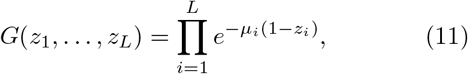

where *µ*_*i*_ is the average number of RNAPs in segment *i* and is given by Eq. (22) in which *η* = *s*_*u*_*/*(*s*_*u*_ + *s*_*b*_) = 1. This generating function leads to a probability distribution that is a product of Poisson distributions,

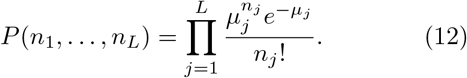

Consequently there are no correlations between segments. Models of this type have been used to describe mRNA senescence [23] and splicing [24].

#### 3. Deterministic elongation with mature RNA production

We can extend this model to include mature RNA production by another stage after transcription termination. The model now has *L* + 1 segments, whereby the first *L* segments describe RNAP dynamics and the last segment counts the number of mature RNA *m*. We further assume uniform elongation rates in segments *i* = 1, …, *L* so that *k*_*el,i*_ = *k*_el_. At the last segment, the mature RNA is degraded at rate *d*_*m*_, which in our notation is equivalent of saying that *d*_*m*_ ≡ *k*_el,*L*+1_ + *d*.

For this model the distribution of mature RNA, *P*(*m*), has been found analytically [16] in the limit in which the elongation is deterministic, i.e when *d*_*m*_*/k*_el_ → 0 and *L* → ∞ but keeping the transcription time *T* = *L/k*_el_ fixed. In this limit, the distribution of the mature RNA is the same as in the telegraph model; hence this limit is a more formal way of deriving the telegraph model from a detailed model of RNAP dynamics.

## III. EXACT MOMENTS OF THE RNAP DISTRIBUTION *P*

In this Section we consider the first three moments of the RNAP distribution *P* in the steady state. We will need these moments later in Section IV, where we derive an analytical expression for the RNAP distribution using mean-field theory.

We begin by deriving recurrence relations for the first three moments, which we solve analytically in the uniform case (all *k*_el,*i*_ are equal to *k*_el_). In the non-uniform case (arbitrary values of *k*_el,*i*_ in each segment) we obtain the first moments analytically and the second and third moments numerically.

In the former case, the first two moments have been previously derived in Ref. [16]. Here for the uniform case we present a new derivation using lattice paths which has the advantage that it can be extended to higher moments, as we demonstrate by computing the third moment.

We first make the following change of variables,

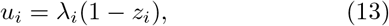

where *λ*_*i*_ is given by

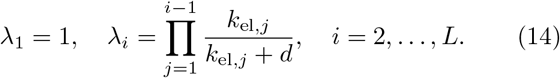

The equations for *G*_on_ and *G*_off_ now take a simpler form,

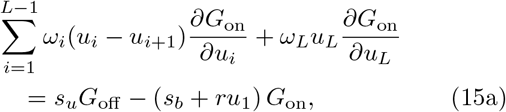

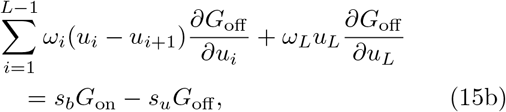

where we have introduced *ω*_*i*_ defined as

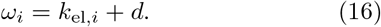

The recurrence relations for the moments of *P* can be found by taking partial derivatives of the equations above and setting *u*_1_ = … = *u*_*L*_ = 0.

### A. First moments of *P*

We introduce the following notation,

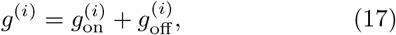

where

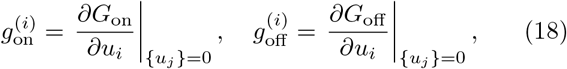

and {*u*_*j*_} = 0 means that *u*_1_ = …= *u*_*L*_ = 0. In order to find *g* we add Eqs. (15a) and (15b) together, then take a partial derivative with respect to *u*_*i*_ and set all *u*_1_ =…= *u*_*L*_ = 0. The resulting recurrence relation for *g*^(*i*)^ reads

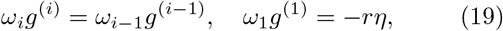

where *η* is the fraction of the time that the promoter spends in the on state,

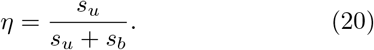

From here we get

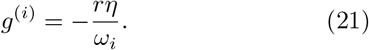

The first moment of *P* in segment *i* is thus given by

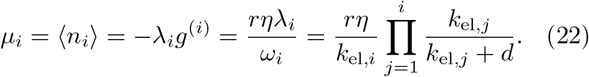

For uniform rates with no detachment, the mean is same across the gene. Otherwise the mean varies across the gene, for example if the elongation rates are non-uniform with no detachment or if the elongation rates are uniform with non-zero detachment

We also write down an analytical expression for 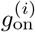 because we will need it later for computing the second moments of *P*,

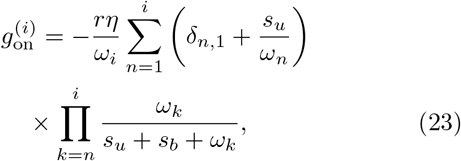

where *δ*_*i,j*_ is the Kronecker delta function. In the case of uniform elongation rates, the expression for 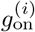 takes a simpler form:

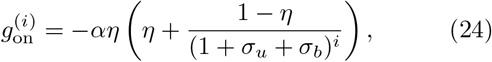

where the rescaled parameters *α, σ*_*u*_ and *σ*_*b*_ are defined in Eq. (9).

### B. Second moments of *P*

In order to find the second moments of *P* we define

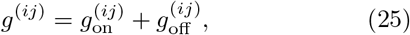

where

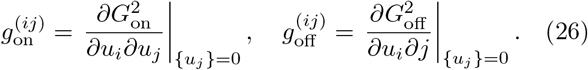

Again, we add Eqs. (15a) and (15b) together, then take partial derivatives with respect to *u*_*i*_ and *u*_*j*_ and set *u*_1_ = …= *u*_*L*_ = 0, yielding the following recurrence relations for *g*^(*ij*)^,

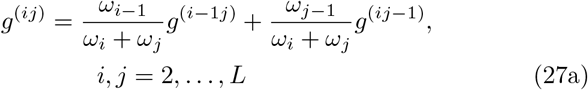

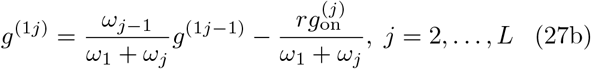

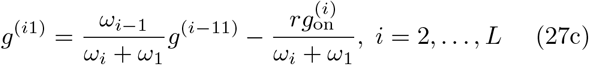

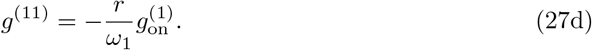

A closed-form expression for *g*^(*ij*)^ can be found in the case of uniform elongation rates using lattice path methods as we show below. In the non-uniform case it is possible to solve the recurrence relations directly (as lattice path methods cannot be used), but it is simpler to solve them numerically. Once we find *g*^(*ij*)^, the covariance cov(*n*_*i*_, *n*_*j*_) can be found from

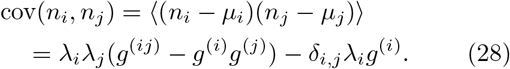

In the case of uniform elongation rates *ω*_*i*_ are all equal and the coefficients in Eqs. (27) become all equal to 1*/*2. We can think of Eqs. (27) in terms of two-dimensional lattice paths with unit steps (*m, n*) → (*m* −1, *n*) and (*m, n*) → (*m, n* − 1). We are interested in the paths that start at (*i, j*) and end at one of the boundary points, (1, *n*) or (*m*, 1), for 1 ≤ *m* ≤ *i* and 1 ≤ *n* ≤ *j*. Let us say we start at (*i, j*) and end at (*m*, 1) for 1 ≤ *m* ≤ *i*. Each step we make is weighted by factor 1*/*2, until we reach the end point at (*m*, 1), which is weighted by 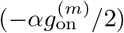, where *α* = *r/*(*k*_el_ + *d*). Similarly, the end point at (1, *n*) for 1 ≤ *n* ≤ *j* is weighted by 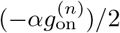, whereas the end point at (1, 1) is weighted by 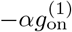.

One such lattice path is presented in Fig. 2 with the start point at (5, 4) and the end point at (2, 1). The total weight of this path is 1*/*2^6^—the length of the path is 6— multiplied by the weight at the end point, 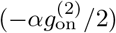, which gives

**FIG. 2.**
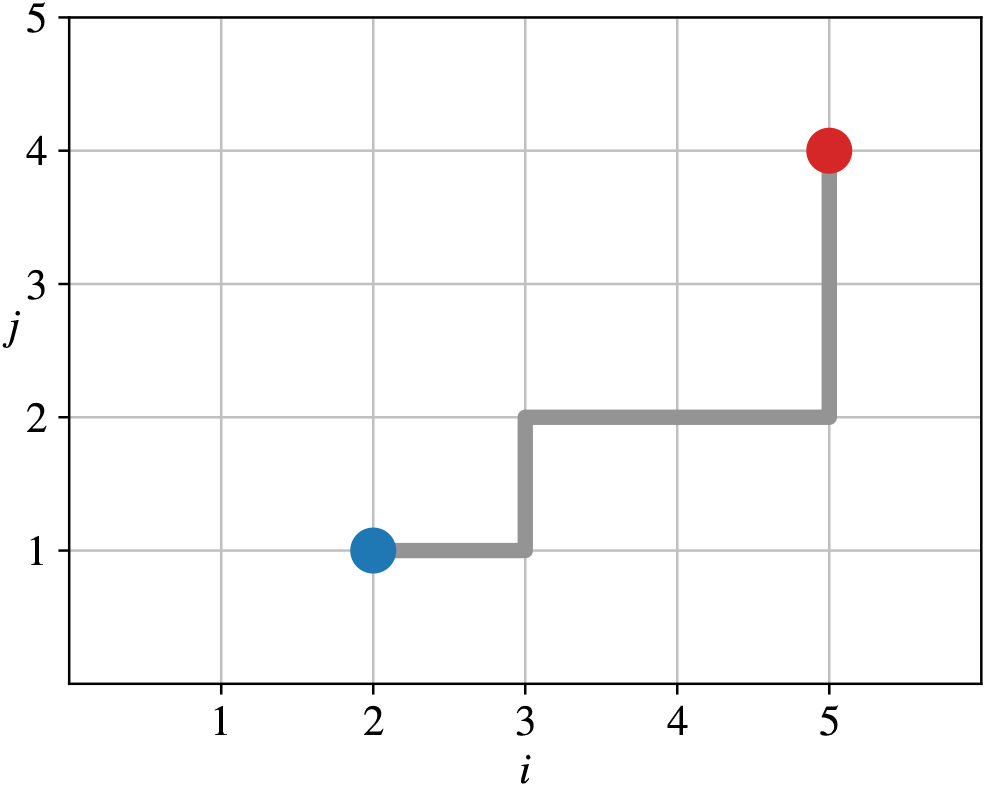
A lattice path from point (*i, j*) = (5, 4) (red) to point (2, 1) (blue). The contribution from this path to *g*^(54)^ is 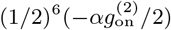.

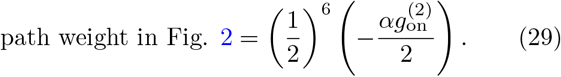

Because all paths between two given points on the lattice carry the same weight, we can simply count them, multiply their weight by the weight at the boundary and then sum over all boundary points. This simplification is not possible in the case of non-uniform elongation rates because paths between two given points on the lattice will in general all have different weights.

In order to count the paths between two fixed points, we denote by *S* moving south and by *W* moving west. We can then describe a path of length *n* as an *n*-letter word consisting of letters S and W. For example, the path in Fig. 2 can be written as SSWWSWW.

Next, we count all paths from a given point (*i, j*) to the boundary point (1, *n*) for each *n* = 2, …, *j*. To get to this point we need to make *i*−1 steps to the south and *j* − *n* steps to the west.(The tota)l length of this path is *i*−1+*j*−*n* and there are 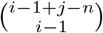 such paths. We repeat this procedure for all paths from (*i, j*) to the boundary points (*m*, 1) for each *m* = 2, …, *i*. Finally, we count all paths from (*i, j*) to (1, 1). After rearranging terms, the total contribution from all paths can be written as

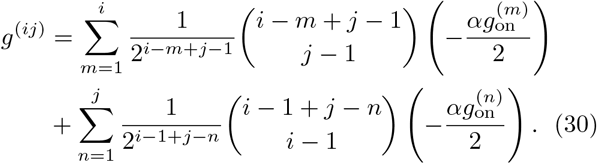

Inserting the expression for 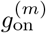 from Eq. (24) into (30) we get a closed-form expression for the second moment of *P* (the covariance)

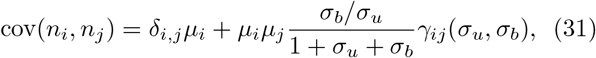

where *γ*_*i,j*_(*σ*_*u*_, *σ*_*b*_)[25] is given by

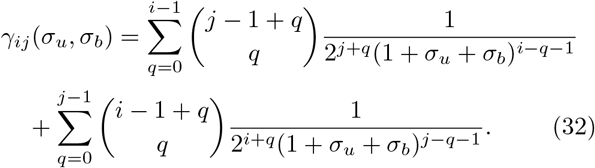

The derivation of the second moments conditioned on the promoter being in the on state can be done is a similar fashion. We omit here the details and state only the final result:

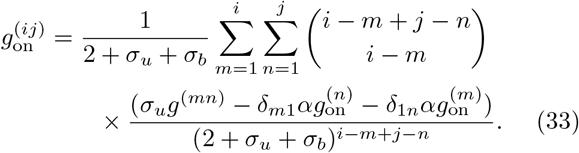

It is interesting that while RNAPs in this model are non-interacting (dynamics of a single RNAP does not depend on the presence of other RNAPs), yet the model exhibits long-range correlations in RNAP numbers between segments. These correlations are entirely due to promoter switching, because the model with constitutive gene expression (*s*_*b*_ = 0) has covariance cov(*n*_*i*_, *n*_*j*_) = 0 for *i* ≠ *j*, see Section II C.

### C. Third moments of *P*

In order to compute third moments pf *P*, we consider third-order partial derivatives of *G*_on_ and of *G*_off_,

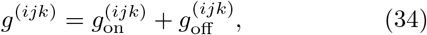

where

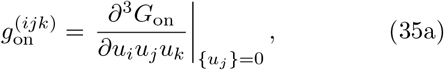

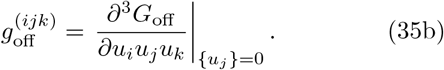

The recipe to obtain the recurrence relation for *g*^(*ijk*)^ is the same as before. We first add Eqs. (15a) and (15b) together, then take the partial derivatives with respect to *u*_*i*_, *u*_*j*_ and *u*_*k*_ and finally set *u*_1_ = … = *u*_*L*_ = 0. The resulting recurrence equations are given by

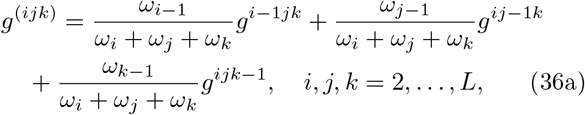

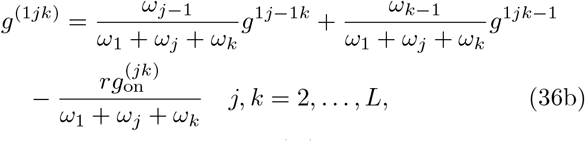

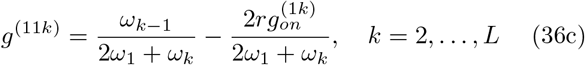

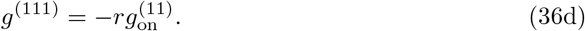

Other equations, for example for *g*^(*i*1*k*)^, can be obtained from these equations using that fact *g*^(*ijk*)^ is invariant under the permutation of indices *i, j, k*. Eqs. (36a) can be solved analytically in the case of uniform elongation rates using lattice paths, or numerically in the non-uniform case. In the former case we consider three-dimensional lattice paths that start at (*i, j, k*) and end at one of the boundary points (1, *n, o*), (*m*, 1, *o*) or (*m, n*, 1), where 1 ≤ *m* ≤ *i*, 1 ≤ *n* ≤ *j* and 1 ≤ *o* ≤ *k*. Instead of 1*/*2, each step is now weighted by 1*/*3. If the end point is, say (1, *n, o*), where *n, o* ≠ 1, then the weight of that point is 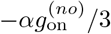, where 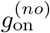 is the second moment conditioned on the promoter being in the on state. If the end point is (1, 1, *o*) for *o* ≠ 1, then the weight is 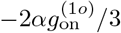. If *o* = 1 then the weight is 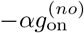. The final result for *g*^(*ijk*)^ is

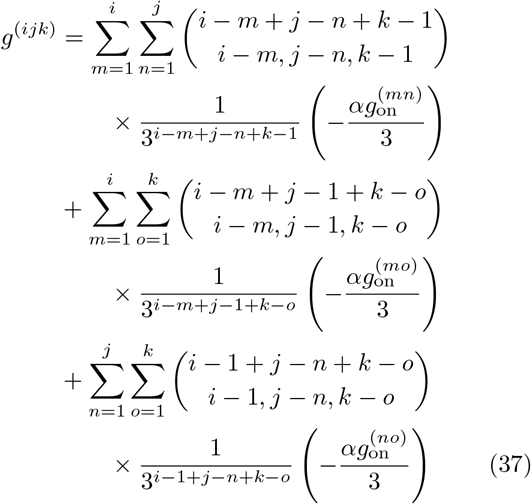

where 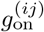 is given by Eq. (33).

Of particular importance is the third standardised central moment in segment *i*, which is given by

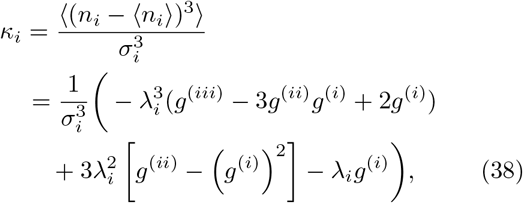

where *g*^(*i*)^, *g*^(*ij*)^ and *g*^(*ijk*)^ are given by Eqs. (21), (30) and (37), respectively, and *σ*_*i*_ is the standard deviation of *n*_*i*_,

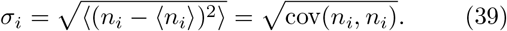

The third standardised central moment or skewness is useful for understanding the shape of the distribution, in particular its (a)symmetry. We will use this expression later in Section IV.

## IV. THE RNAP DISTRIBUTION IN THE MEAN-FIELD APPROXIMATION

As we mentioned earlier, solving the system of partial differential equations for *G*_on_ and *G*_off_ for arbitrary number of segments *L* is a difficult problem. In order to make progress, we focus on the marginal distribution of *n*_*i*_ in the steady state defined as

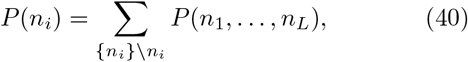

where the summation goes over all values of variables *n*_1_, …, *n*_*L*_ except for the variable *n*_*i*_ which is kept fixed. In this section we provide an analytical expression for *P*(*n*_*i*_) in the mean-field approximation.

### A. Naive mean-field approximation

We first consider an approximation that we call the naive mean-field approximation (NMF). This type of approximation is well known in statistical physics and often gives satisfactory results in the absence of long-range correlations (for example, away from phase transitions). However, we will show that for this model the naive mean-field approximation does not always work, and later we find an improved mean-field approximation that shows excellent agreement with the exact results from stochastic simulations.

The main idea of the naive mean-field approximation is to find an effective master equation for the marginal distribution *P*(*n*_*i*_) that is decoupled from fluctuations in segment *I* − 1 in which case we can solve it analytically. We do this by summing Eqs. (4a) and (4b) over all *n*_*j*_ for all *j* ≠ *i*. Many terms cancel in the summation and we are left with master equations for *P*_on_(*n*_*i*_) and *P*_off_(*n*_*i*_). In each master equation there will be only one term that couples segment *i* with *i* −1, and that is the term that describes elongation from *i* −1 to *i*. For example, in the master equation for *P*_on_, this term is given by

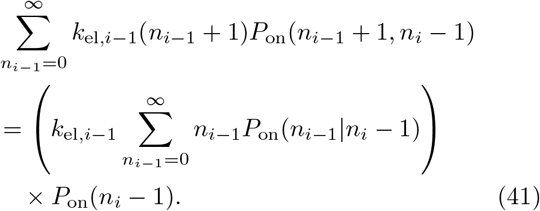

The expression in the parentheses is the effective rate at which new RNAPs are added to segment *i* from segment *i* − 1. The problem is of course that we cannot compute this rate unless we compute the conditional distribution *P*_on_(*n*_*i*−1_ |*n*_*i*_ − 1). The naive mean-field approximation amounts to replacing

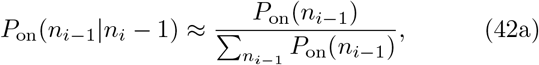

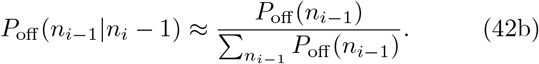

In other words we ignore correlations between segments *i* − 1 and *i*, except for the fact that both segments have the same promoter state. The effective rates *p*_*i*_ and *q*_*i*_ at which new RNAPs are added to segment *i* in the on and off state are thus given by

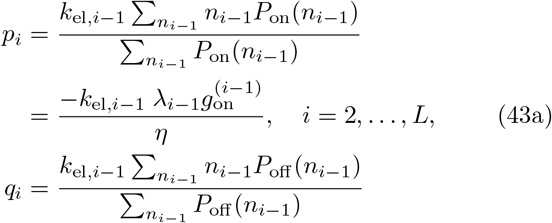

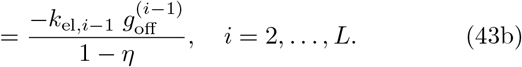

The effective model that governs the stochastic dynamics of *n*_*i*_ in the naive mean-field approximation is equivalent to the telegraph model for RNA production [5] in which transcription rates in the on and off states are *p*_*i*_ and *q*_*i*_, respectively, and the mRNA degradation rate is *k*_el,*i*_ + *d*. The reactions for this model with their corresponding rates are given by

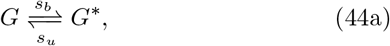

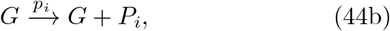

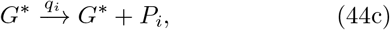

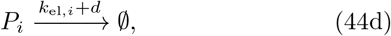

where *G* and *G*^∗^ represent the on and off states of the gene and *P*_*i*_ is the species associated with the *i*^*th*^ segment (see scheme (1)). We note that the telegraph model in which transcription is allowed from both on and off states is also known as the leaky telegraph model [11].

The telegraph model described by reactions (44a)-(44d) can be solved exactly and the generating function reads

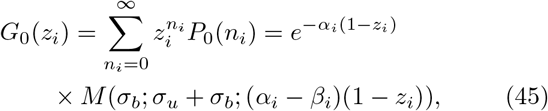

where *α*_*i*_, *β*_*i*_, *σ*_*u,i*_ and *σ*_*b,i*_ are defined as

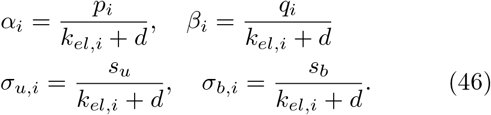

The probability distribution *P*(*n*_*i*_) in the naive meanfield approximation reads

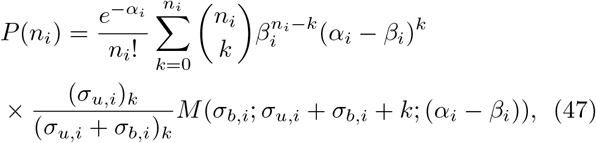

and the mean and the variance are given by, respectively,

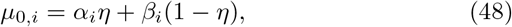

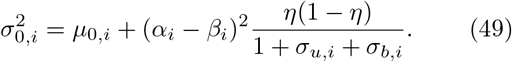

We can check that the choice of *p*_*i*_ and *q*_*i*_ in Eqs. (43a) and (43b) ensures that *µ*_0,*i*_ = *µ*_*i*_, i.e. the mean of *n*_*i*_ computed in Eq. (22) is the same as the mean in the naive mean-field approximation. However the value of the variance is generally different from the exact value.

In the case of uniform elongation rate the expression for the variance simplifies to

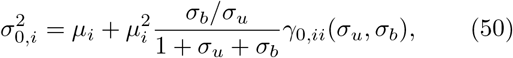

where *γ*_0,*ii*_(*σ*_*u*_, *σ*_*b*_) is given by

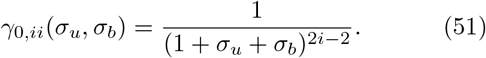

The difference between the exact variance in Eq. (31) and Eq. (50) is in the factor *γ*_*ii*_ for *i* ≥ 2 (the naive mean-field approximation is exact in segment 1). In general we find that *γ*_0,*ii*_ *< γ*_*ii*_ for any *i* = 2, …, *L* for *σ*_*u*_ ≠0 and *σ*_*b*_ ≠ 0, whereas *γ*_0,*ii*_ = *γ*_*ii*_ = 1 for *σ*_*u*_ = *σ*_*b*_ = 0. The naive mean-field approximation underestimates the exact variance, which is expected given that we ignored correlations. When *σ*_*b*_ = 0 (constitutive gene expression), there are no correlations, the naive mean-field approximation becomes exact and the resulting distribution is Poisson, see Eq. (12).

In general, the naive mean-field approximation is valid if the correlations between *n*_*i*−1_ and *n*_*i*_ are small. Fortunately, for this model we can compute these correlations exactly from the covariance cov(*n*_*i*−1_, *n*_*i*_) in Eq. (28). In particular we can compute the correlation coefficient defined as

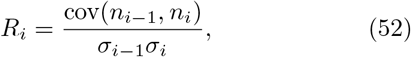

where *σ*_*i*_ is the standard deviation of *n*_*i*_, see Eq. (39). The naive mean-field is valid provided *R*_*i*_ ≈ 0, i.e. when *n*_*i*−1_ and *n*_*i*_ are not correlated. Computing *R*_*i*_ thus gives us a direct method to check the validity of the naive mean-field theory.

In Fig. 3 we compare RNAP number distribution in the naive mean-field approximation with the exact distribution obtained from stochastic simulations, for two segments on the gene, *i* = 2 and *i* = *L* = 30. In the case of constitutive gene expression (Fig. 3(a)) we find an excellent agreement between the distributions, in accordance with the fact *R*_2_ = 1.5 · 10^−3^ and *R*_30_ = 10^−4^. In the case of bursty expression the agreement is not so good for *i* = 2, but improves for *i* = *L* (Fig. 3(b)). Again, this can be explained by stronger correlations at the start of the gene than at the end, *R*_2_ = 0.35 and *R*_30_ = 0.18. The disagreement between the distributions is strongest in the bimodal and nearly bimodal cases (Figs. 3(c) and (d)). In those cases the correlations are the strongest: we find *R*_2_ = 0.74 and *R*_30_ = 0.58 for the bimodal and *R*_2_ = 0.76 and *R*_30_ = 0.59 for the nearly bimodal distributions.

**FIG. 3.**
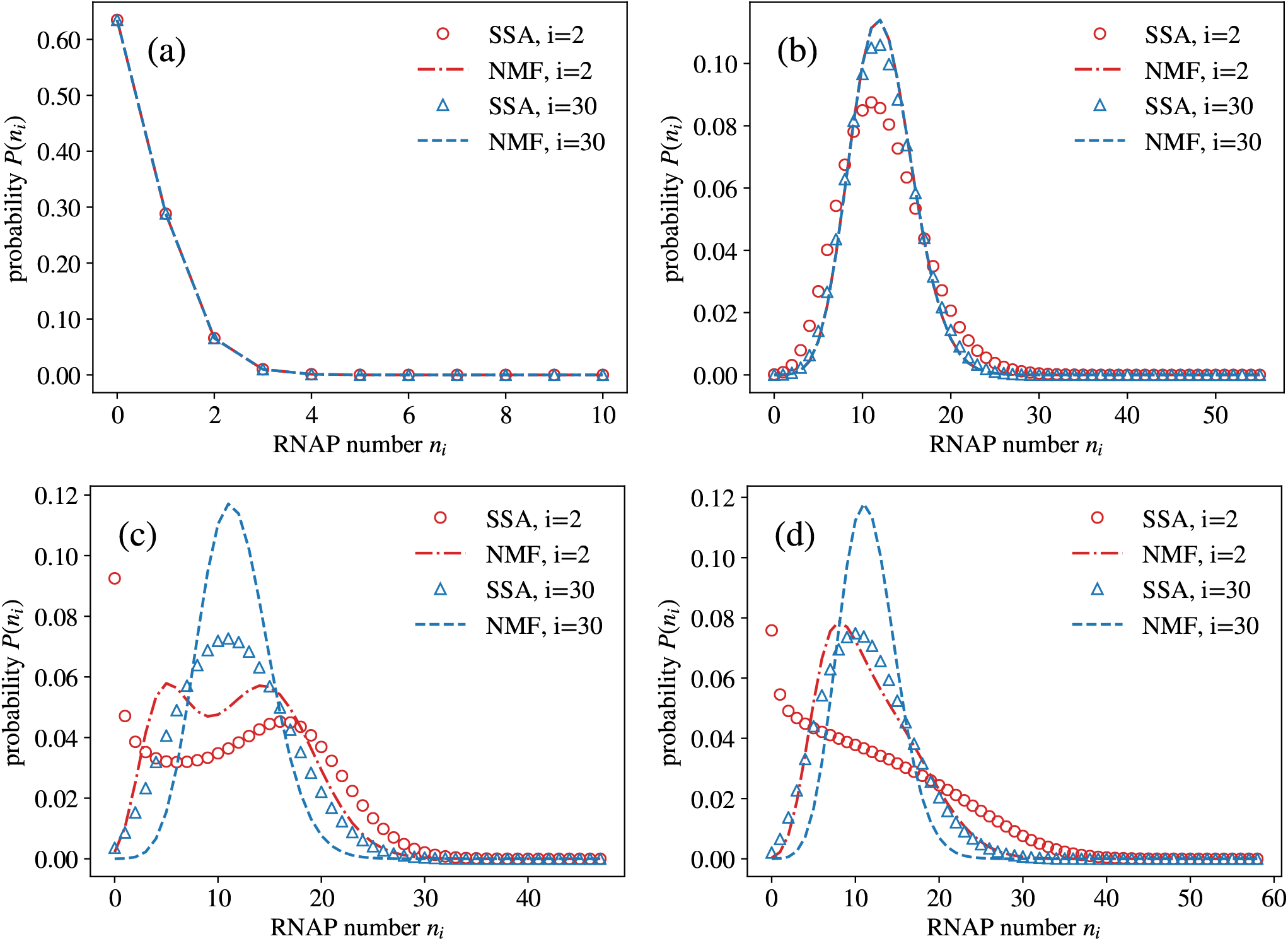
RNAP number distribution in segments *i* = 2 and *i* = *L* = 30 in the naive mean-field approximation (NMF; the distribution is given by Eq. (47)), compared to the exact distribution obtained by using the stochastic simulation algorithm (SSA). Elongation rates are assumed to be uniform (*k*_el,*i*_ = *k*_el_). (a) Constitutive expression: *r* = 0.5, *k*_el_ = 1, *d* = 0, *s*_*u*_ = 10, *s*_*b*_ = 1. (b) Bursty expression: *r* = 100, *k*_el_ = 1, *d* = 0, *s*_*u*_ = 7, *s*_*b*_ = 50. (c) Bimodal distribution: *r* = 20, *k*_el_ = 1, *d* = 0, *s*_*u*_ = 0.35, *s*_*b*_ = 0.25. (d) Nearly bimodal distribution: *r* = 30, *k*_el_ = 1, *d* = 0, *s*_*u*_ = 0.5, *s*_*b*_ = 0.8.

### B. Improved mean-field approximation

We saw in the previous section that the naive meanfield approximation is equivalent to a telegraph model that produces exact mean but wrong variance. In this section we consider a telegraph model with general rates,

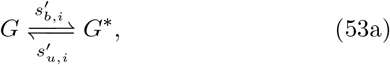

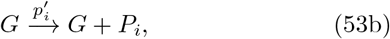

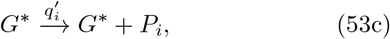

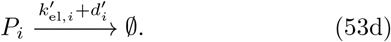

We want to find rates 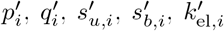 and 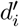 such that the mean 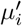 and variance 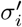 of the telegraph model described by Eqs. (53a)-(53d) are both matched to their exact values *µ*_*i*_ in Eq. (22) and 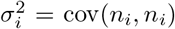 in Eq. (28), respectively. In particular, we want to solve

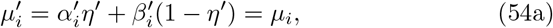

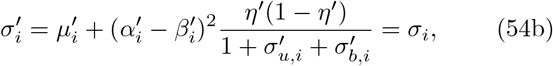

where 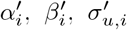 and 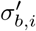 are obtained as before by rescaling 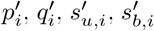 by 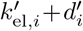, and 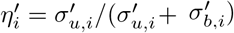.

Obviously we cannot find unique rates that solve Eqs. (54a)-(54b) because the problem is overdetermined. We therefore keep the elongation rate 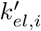 and the detachment rate 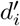 the same as in the original model,

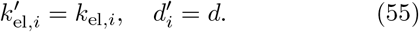

That leaves us with four parameters to adjust by solving two equations, Eqs. (54a) and (54b). We further require that the fraction of time the gene spends in the on state, 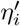, is preserved,

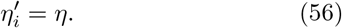

We consider three options for the remaining parameters:

- Option 1: We set 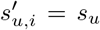 and 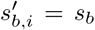 so that Eq. (56) is automatically satisfied. We then adjust the effective transcription rates 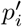 and 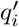 to match the mean and variance. This is a leaky telegraph model with the same gene switching rates as in the original model.
- Option 2: We set 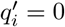 and adjust 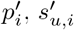 and 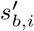 by solving Eqs. (54a), (54b) and (56). This is a non-leaky telegraph model with effective, segment-dependent gene switching rates.
- Option 3: We adjust all four rates 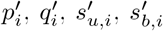 by solving Eqs. (54a), (54b) and (56). Additionally we require that the skewness predicted by the telegraph model, Eq. (59), is matched to the exact skewness computed in Eq. (38). This is a leaky telegraph model with effective, segment-dependent gene switching rates.

Surprisingly, option 1 does not lead to a noticeable improvement over the naive mean-field approximation, and we discard it. In contrast, option 2 significantly improves the naive mean-field approximation in all four cases: the constitutive, bursty, bimodal and nearly bimodal, as we demonstrate below. Interestingly, option 3 does not provide noticable improvement over option 2. We will show later that that is because option 2 predicts skewness that is very close to the exact one. Since option 2 has a simpler expression for the probability density, we from now on consider only option 2 to which we refer to as the improved mean-field approximation (IMF).

For option 2 the solution to Eqs. (54a), (54b) and (56) with 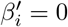 is unique and reads,

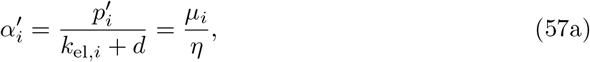

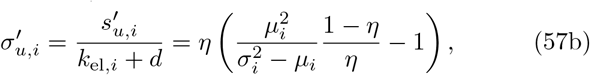

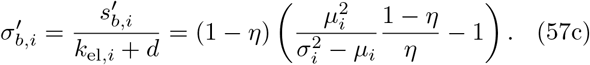

One can show that the effective parameters 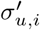 and 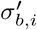 are always positive.

These parameter values can then be used to compute the probability distribution in the improved mean-field approximation (IMF) from the telegraph model,

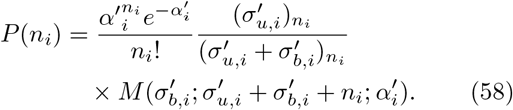

In Fig. 4 we compare the approximate distribution in Eq. (58) to the exact distribution obtained by stochastic simulations using the same model parameters *α, σ*_*u*_ and *σ*_*b*_ as in Fig. 3. We find an excellent agreement in all four cases, the constitutive, bursty, bimodal and nearly bimodal. We argue that option 2 performs significantly better than option 1 because the correlations between segments are solely due to promoter switching. These correlations can be better taken into account by adjusting the effective on and off rates, rather than keeping them the same.

**FIG. 4.**
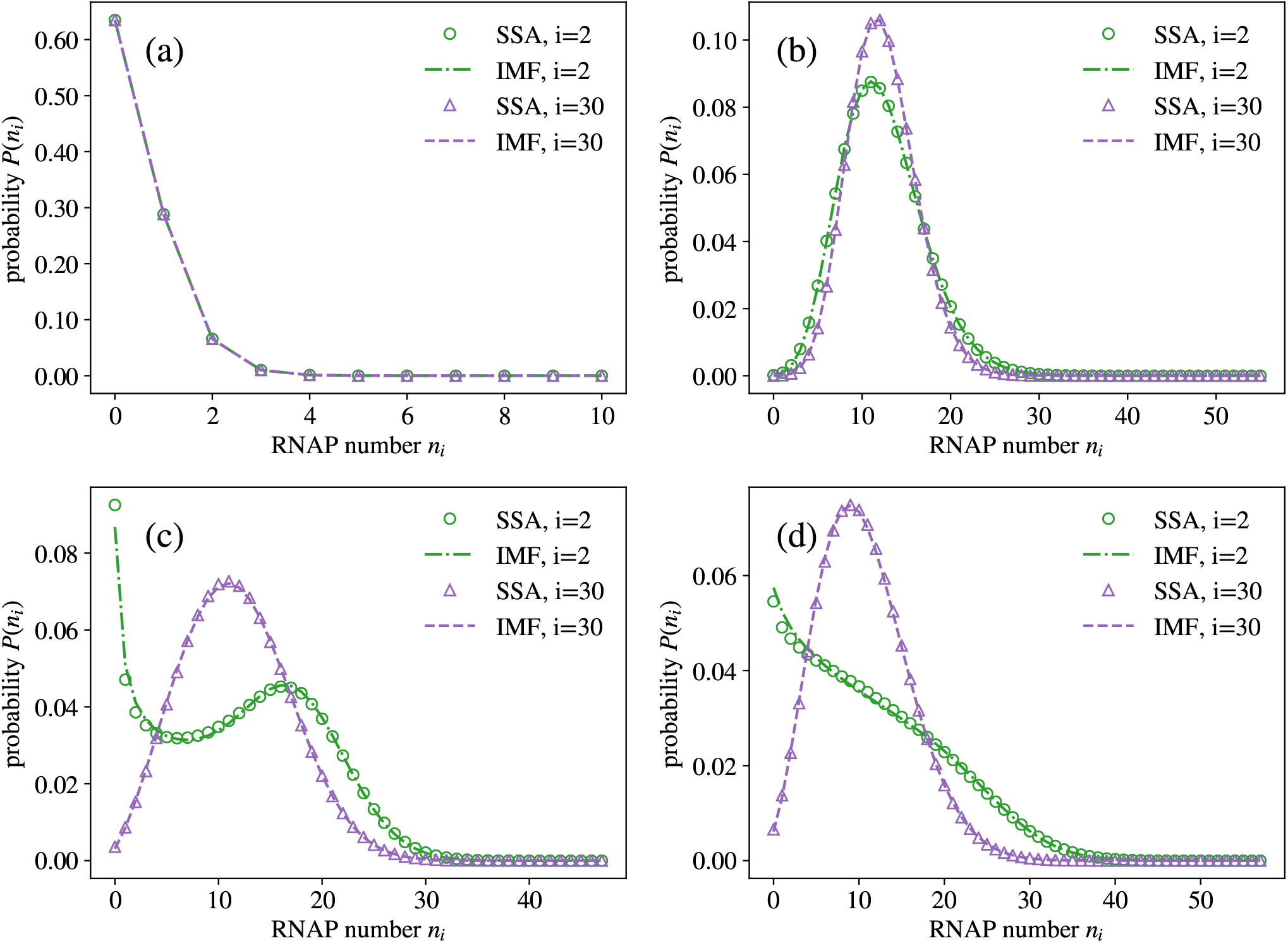
RNAP number distribution in segments *i* = 2 and *i* = *L* = 30 in the improved mean-field approximation (IMF; the distribution is given by Eq. (58)), compared to the exact distribution obtained by the stochastic simulation algorithm (SSA). Model parameters are the same as in Fig. 3. The improved mean-field approximation performs significantly better than the naive mean-field approximation and is accurate even when the number correlations between segments are large. (a) Constitutive expression. (b) Bursty expression. (c) Bimodal distribution. (d) Nearly bimodal distribution.

Because the improved mean-field approximation matches only the first two moments, we wanted to check its accuracy in predicting the third standardised central moment or skewness. To this end we generated 8000 unique values of the parameters *α, σ*_*u*_ and *σ*_*b*_. The values for each parameter were generated uniformly on a logarithmic scale using according to formula 2^*k*^ for integer *k* between −13 and 6. That gave us 20 values for each parameter spanning over five orders of magnitude in the range from 1.2 ·10^−4^ to 64. For each combination we computed the exact skewness *κ*_*i*_ according to Eq. (38), and compared it to the one predicted by the improved mean-field approximation (computed from Eq. (58)),

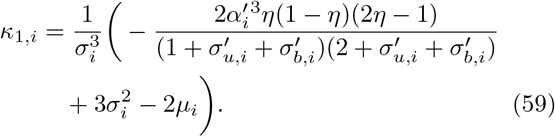

We did this for all segments on the gene, after which we selected the segment with the largest relative error *ϵ* = 100% ·|*κ*_1,*i*_ − *κ*_*i*_ |*/κ*_*i*_. For this segment we further computed the RNAP number distribution in the improved mean-field approximation according to Eq. (58). Depending on the shape, we determined to which of the three categories the distribution corresponds to: unimodal (without an inflection point), bimodal and nearly bimodal (unimodal with an inflection point). Of 8000 distributions, 7190 were unimodal, 487 bimodal and 323 nearly bimodal.

The results comparing *κ*_1,*i*_ versus exact *κ*_*i*_ are presented in Fig. 5. Due to the large range of *κ*_*i*_, we separated the data according to *κ*_*i*_ *<* 10 (Fig. 5(a), linear scale) and *κ*_*i*_ *>* 10 (Fig. 5(b), log scale). Next, we inspected in more detail how the relative error *ϵ* is distributed among the four aforementioned categories. We focused on distributions for which the relative error *ϵ >* 10%. We found 74 such distributions in total, 64 were unimodal and 10 were bimodal. A closer inspection of these distributions revealed that they all had in common skewness *κ*_*i*_ ≈ 0 which caused the relative error to be large. We inspected the distributions with the two largest *ϵ* values and used the Hellinger distance [26] to quantify their similarity compared to the exact distributions obtained by stochastic simulations. Indeed, we found very small Hellinger distances for both distributions, 3.7 ·10^−3^ and 3.3 ·10^−3^, confirming the high accuracy of our improved mean-field approximation.

**FIG. 5.**
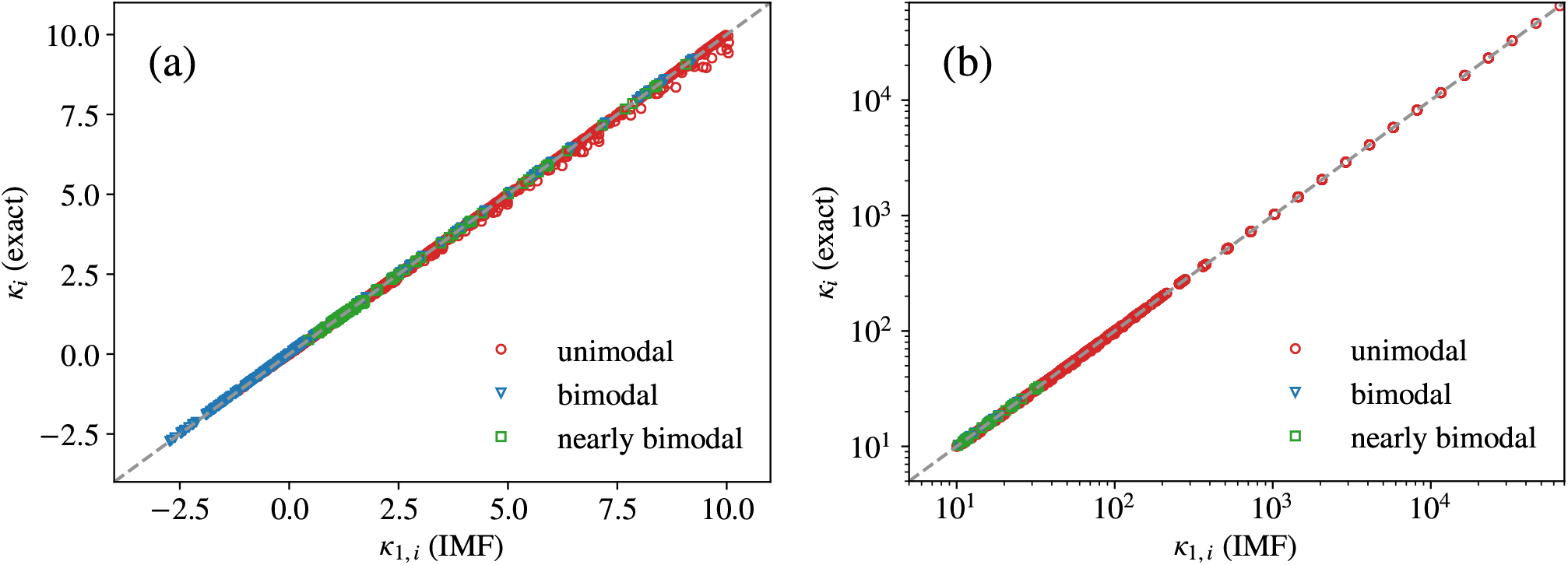
Skewness *κ*_1,*i*_ predicted by the improved mean-field approximation (IMF; see Eq. (59)) compared to the exact value *κ*_*i*_ (given by Eq. (38)) for 8000 combinations of *α, σ*_*u*_ and *σ*_*b*_. Each point (*κ*_1,*i*_, *κ*_*i*_) corresponds to one combination of parameters *α, σ*_*u*_ and *σ*_*b*_. For each combination, the segment *i* with the largest relative error between *κ*_1,*i*_ and *κ*_*i*_ was selected. The results were divided in two sets: (a) is for *κ*_*i*_ *<* 10 (linear scale) and (b) is for *κ*_*i*_ *>* 10 (log scale). The dashed line is for reference only and given by *κ*_1,*i*_ = *κ*_*i*_.

### C. Other applications: nuclear retention and export of mRNA

So far we have analysed the performance of the improved mean-field theory in the context of transcription elongation for which the assumption of uniform elongation seems reasonable. However, in other multistep downstream processes such as splicing or transport of nuclear mRNA to the cytoplasm, we cannot expect the rates of different downstream processing steps to be equal because each step may represent a physically different process.

Here we demonstrate the performance of the improved mean-field theory for the case of non-uniform rates. We consider the model for nuclear mRNA retention developed in Ref. [18]. A reaction scheme for this model with the corresponding rates is given by

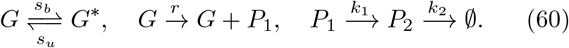

In this model, nuclear mRNA *P*_1_ is produced at rate *r* from a gene in the on state and exported at rate *k*_1_ from the nucleus to the cytoplasm (this is an *L* = 2 version of the reaction scheme (1)). The cytoplasmic mRNA *P*_2_ is degraded at rate *k*_2_. Parameters for this model were quantified experimentally for a set of genes in mouse liver cells in Ref. [18]. For our purposes we will use the following parameters obtained for Mlxipl gene,

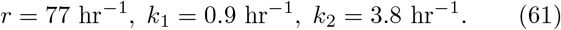

In Ref. [18] only the fraction of active Mlxipl genes *η* = 0.425 was measured but not the absolute on and off rates *s*_*u*_ and *s*_*b*_. In addition the coefficient of variance (CV) for cytoplasmic mRNA was measured to be CV= 0.46. By matching *η* and CV to those predicted by our theory (using the results for the mean and variance), we estimated the switching parameters: *s*_*u*_ = 4.3 hr^−1^ and *s*_*b*_ = 5.7 hr^−1^. In addition to these values we considered two additional values of export rate *k*_1_ that were two times smaller and larger than the experimental value.

Using these parameters we performed stochastic simulations of the model and compared the distributions of *P*_1_ and *P*_2_ with predictions of the improved mean-field theory. The results for nuclear mRNA are presented in Fig. 6(a) and for cytoplasmic mRNA in Fig. 6(b). In all cases we find excellent agreement with the predictions of the improved mean-field theory showing that it is equally accurate for non-equal rates of downstream processing steps.

**FIG. 6.**
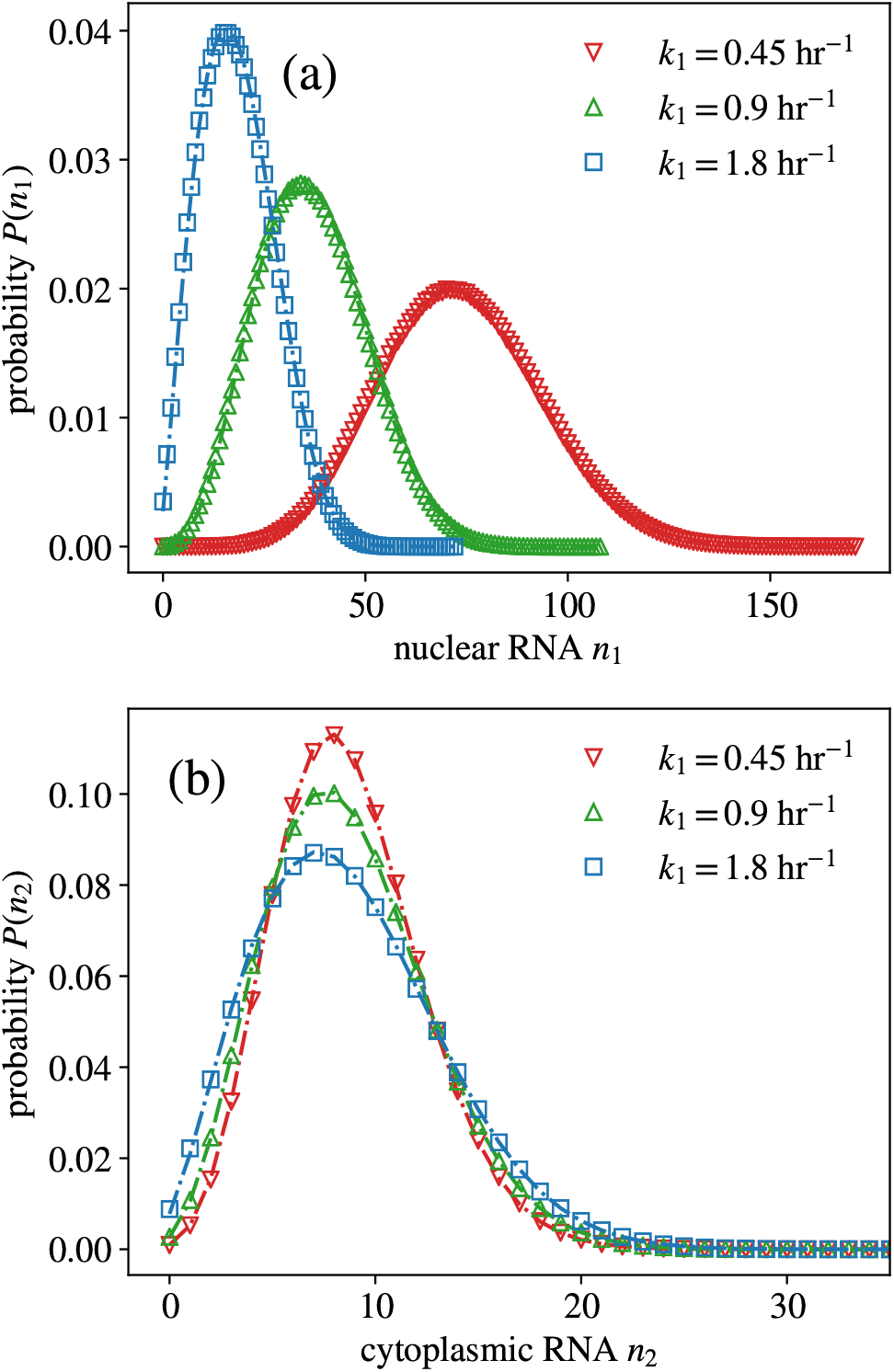
Probability distributions of nuclear and cytoplasmic mRNA numbers in a two-state model of transcription with nuclear retention. Distributions were obtained by stochastic simulations (symbols) and compared to the prediction of the improved mean-field theory (lines). (a) Distribution of nuclear mRNA number *P*(*n*_1_). (b) Distribution of cytoplasmic mRNA number *P*(*n*_2_). The reaction scheme is given by (60) and the parameters are stated in the main text.

### D. Variation of the shape of the distribution with increasing number of downstream processing steps

In Fig. 4(c), we saw how a distribution that is bimodal for small *i*, can become unimodal for large enough *i*. The question we want to address here is whether this observation is a special case or if it generally holds.

Where the model is interpreted as one for RNAP dynamics during transcription, assuming uniform elongation rates across the gene (*k*_el,i_ = *k*_el_) and no premature detachment (*d* = 0), using the exact Eqs. (22) and (31), one can easily deduce that the factor 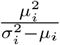 increases monotonically with *i*. Hence from Eqs. (57b) and (57c), it follows that the effective switching rates in the improved mean-field theory increase with distance *i* from the start site. This implies that any bimodality in the RNAP distribution is washed out as *i* increases because bimodality typically manifests due to slow switching between inactive and active states [27, 28].

A difficulty in generalizing this statement to the case of non-identical elongation rates and non-zero premature detachment is that in this case while the recurrence relations for the moments can be solved analytically, the solution is very complex for general number of down-stream processing steps *L*. Instead it is easier to use Mathematica to solve the recurrence relations for *L* = 1, then for *L* = 2, etc and try to see a pattern in the algebraic equations. Using this method we verified that for any integer *L*, the factor 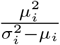 increases monotonically with *i*. Hence by the same arguments as in the previous paragraph, we can generally state that for the general model (1), any bimodality in the distribution of *P*_*i*_ will decrease as *i* increases. Interpreting reaction scheme (1) as a model of the whole mRNA lifecycle, this implies that in a population of identical cells (since our model does not consider cell-to-cell variability), it is easier to observe bimodality in nascent/nuclear mRNA distributions than in cytoplasmic distributions.

The increase of the effective switching rates with *i* has also implications for the inference of the extent of transcriptional bursting [29]. This phenomenon is characterized by bursts of mRNA expression separated by long intervals of no expression, i.e. large mean burst size and low burst frequency. These two parameters are often estimated by fitting the telegraph model to measured distributions of the mRNA copy number [30]. Now it is well known that within the mathematical framework of the (non-leaky) telegraph model of gene expression, the mean burst size is the initiation rate divided by the off switching rate while the burst frequency is the on switching rate. We have shown above that the effective on rate increases with *i* which means that fitting the telegraph model to mRNA data primarily described by the *i*^*th*^ stage of the life cycle will necessarily overestimate the true burst frequency. The effective mean burst size is obtained by dividing Eq. (57a) by Eq. (57c). While the latter equation increases with *i*, the former equation can be independent or increase or decrease with *i*. Hence fitting the telegraph model to mRNA data primarily described by the *i*^*th*^ stage of the life cycle can under or overestimate the true burst size.

These results show that noise due to downstream processing steps can have a significant effect on the estimation of transcriptional parameters; this makes the case that for the accurate estimation of these parameters, the use of nascent mRNA data (which would correspond to small *i* in our model) is preferable to the use of cytoplasmic or whole cell mRNA data [31].

## V. SUMMARY AND DISCUSSION

In this paper, we have devised a simple but very effective approximation to the steady-state distribution of the reaction scheme (1). The main idea behind our work was to approximate the dynamics of *P*_*i*_ in this model by that of an effective (non-leaky) telegraph model:

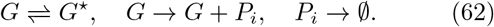

In the improved mean-field approximation, the effective rates of the latter are deduced by matching the first two moments of *P*_*i*_ in the full and reduced models such that the ratio of the switching rates is the same in both models (but the absolute rates are not the same). This procedure is possible because the moments of both models are known exactly due to the linearity of their propensities. We have shown that this automatically means that the third-moments of both models are very close to each other, which suggests that the distributions of both models are also very close. By an extensive search over parameter space, we verified that the steady-state distribution of the reduced model distribution (which is known in closed-form) was very close to that obtained from stochastic simulations of the full model – the maximum Hellinger distance between the two distributions is of the order of 10^−3^ and in fact in all cases one could not easily distinguish by eye any difference between the two.

The study by Vastola et al [17] is the only other study of the steady-state distribution of (1) that we are aware of in the literature. In this paper, the same model is considered but there is no premature degradation and there is a non-zero transcription rate from the off state *G*^∗^. The model is studied in the context of transcription coupled to a switching gene with multiple downstream splicing steps. In this case, *P*_*i*_ can be interpreted as the number of mRNA molecules after *i −* 1 splicing steps. The paper use tools inspired by quantum mechanics, including ladder operators and Feynman-like diagrams, to write a formula for the joint distribution in terms of an infinite sum. This cannot be generally written in closed form. In particular the solution is in powers of the difference between the transcription rates of the active and inactive states, and since this is not typically small, one must compute a large number of series terms in order to closely approximate the correct answer. This is a disadvantage compared to our method which leads to a simple closed-form solution that is easy to evaluate and numerically stable. The disadvantage of our method relative to that of Vastola et al. is that it is not based on a systematic derivation; however as we have shown our method leads to an impressively accurate approximation of the full model over all parameter space. Another work related to our study is that by Bokes and Singh [32] which studied a one state gene system where mRNA molecules are produced in bursts (with a size that is distributed according to the geometric distribution) and then they are exported to the nucleus. This is likely [33] a special case of our model, i.e. the bursty limit where *r* and *s*_*b*_ → ∞ at constant *r/s*_*b*_ for two segments *L* = 2. The authors find an exact solution to this special case; the marginal distribution of mRNA is found to be always unimodal. The advantage of our approximation is that it is valid not just in the bursty limit but all across parameter space, thus being able to capture the variation of the modality of the distribution through the mRNA lifecycle (all three types of distributions – unimodal, bimodal and nearly bimodal – have been measured [34–36]).

Finally we note that our present model could be improved to include further biological realism e.g. an arbitrary number of gene states and time-dependent switching rates. The latter is important since the presence of additional states is sometimes needed to explain the non-exponential duration of the off state for mammalian genes [37]; indeed a recent paper [38] found that a model equivalent to our reaction scheme (1) but with at most three gene states was sufficient to explain the general features of transcription and pervasive stochastic splice site selection. The extension of our model to consider time-dependent switching rates is also important to describe for example the activation of a promoter by a time-dependent transcriptional factor signal e.g. the identities and intensities of different stresses in budding yeast are transmitted by modulation of the amplitude, duration or frequency of nuclear translocation of the general stress response transcription factor Msn2 [39]. The modification of our novel mean-field approach to describe these more complex scenarios will be reported in a separate paper.

## ACKNOWLEDGMENTS

We are grateful to Tatiana Filatova for critical reading of an earlier version of this manuscript. This work was supported by a Leverhulme Trust research award (RPG-2020-327).

